# Crossing enhanced and high fidelity SpCas9 nucleases to optimize specificity and cleavage

**DOI:** 10.1101/187898

**Authors:** Péter István Kulcsér, András Tálas, Krisztina Huszár, Zoltán Ligeti, Eszter Tóth, Nóra Weinhardt, Elfrieda Fodor, Ervin Welker

## Abstract

**Background:** The propensity for off-target activity of Streptococcus pyogenes Cas9 (SpCas9) has been considerably decreased by rationally engineered variants with increased fidelity (eSpCas9; SpCas9-HF1). However, a subset of targets still generate considerable off-target effects. To deal specifically with these targets, we generated new "Highly enhanced Fidelity" nuclease variants (HeFSpCas9s) containing mutations from both eSpCas9 and SpCas9-HF1 and examined these improved nuclease variants side-by-side, to decipher the factors that affect their specificities and to determine the optimal nuclease for applications sensitive to off-target effects.

**Results:** These three increased-fidelity nucleases can routinely be used only with perfectly matching 20 nucleotide-long spacers; a matching 5' G extension being more detrimental to their activities than a mismatching one. HeFSpCas9s exhibit substantially improved specificity specifically for those targets for which eSpCas9 and SpCas9-HF1 have higher off-target propensity. There is also a ranking among the targets by their cleavability and off-target effects manifested by the increased fidelity nucleases. Furthermore, we show that the mutations in these variants may diminish the cleavage, but not the DNA-binding, of SpCas9s.

**Conclusions:** No single nuclease variant shows generally superior fidelity; instead, for highest specificity cleavage, each target needs to be matched with an appropriate high fidelity nuclease. We provide here a framework for generating new nuclease variants for targets that currently have no matching optimal nuclease, and offer a simple mean for identifying the optimal nuclease for targets in the absence of accurate target-ranking prediction tools.

## Background

Cas9 proteins are RNA-guided endonucleases that can be directed to cleave a chosen DNA sequence [1-4]. This process requires complementarity between the Cas9-associated single guide RNA (sgRNA) and the target site in addition to the presence of a short protospacer-adjacent motif (PAM) at the 3'-end of the target [5-13]. Although it has been demonstrated that the *Streptococcus pyogenes* Cas9 (SpCas9) nuclease can be used for genome engineering, its widespread use has been limited by its off-target activity; i.e. the nuclease also cleaves targets that show limited, imperfect complementarities with the associated sgRNA [14-21]. The off-target sequences are difficult to predict and have been shown to contain up to 5-6 base mismatches [22, 23], a property that may interfere with many research applications as well as using this nuclease for therapeutic purposes. Much effort has been devoted to circumvent these confounding effects of the nuclease, such as reducing the amount of active Cas9 in the cell [14, 24, 25], using truncated sgRNAs that bear shortened regions of target site complementarity [15], engineering SpCas9 mutants [26], using paired SpCas9 nickases [27, 28] or using pairs of catalytically inactive SpCas9 fused to a non-specific FokI nuclease domain [29-31]. Recently, attempts to use structure-guided engineering of SpCas9 to reduce its off-target activities have been reported: the enhanced SpCas9 [eSpCas9(1.1), K848A/K1003A/R1060A] [32], hereinafter referred to as eSpCas9, was developed to decrease the affinity of the protein for the non-target DNA strand hence increasing the strand’s propensity for reinvading the RNA-DNA hybrid helix and therefore, decreasing the stability of mismatch-containing helices. By contrast, mutations contained in the high fidelity Cas9 (SpCas9-HF1, N497A/R661A/Q695A/Q926A) [33] weaken the interactions of the protein with the target DNA strand aimed at decreasing the energetics of the SpCas9–sgRNA complex so that it retains a robust on-target activity but has a diminished ability to cleave mismatched off-target sites. Both mutants exhibited considerably reduced off-target effects when assessed by unbiased whole-genome off-target analysis: their cleavage activity toward off-targets with multiple mismatches were almost completely eliminated, although some off-targets, mainly with single-base mismatches, were found.

However, a subset of targets, referred to as atypical, with repetitive or homopolymeric sequences were still cleaved with considerable off-target effects [33]. While these results are very encouraging, it is difficult to decide which SpCas9 variant is superior for applications where the avoidance of off-target activity is of paramount interest, because they were characterized in differing experimental setups: exploiting different sets of targets in different (either U2OS or HEK) cells and employing different methods (GUIDE-seq [18] versus BLESS [34]) to assess their genome-wide specificity.

Here, we generate new variants (“Highly enhanced Fidelity” or HeFSpCas9s) of SpCas9, containing combinations of mutations from both eSpCas9 and SpCas9-HF1, that show higher fidelity specifically with respect to those targets for which eSpCas9 and SpCas9-HF1 exhibit a higher off-target propensity. Furthermore, we directly compare these highly improved nucleases in the same system to understand the factors that affect their specificity and to help select for which off-target sensitive applications each would be most suitable.

## Results

In order to facilitate a thorough comparison of the nuclease variants, we subcloned the wild type and mutant nucleases (eSpCas9, SpCas9-HF1 and the mutants developed in this study, HeFSpCas9 and later HeFm1-and HeFm2SpCas9) into the same plasmid backbone and tailored them to have identical NLS and FLAG tags at their termini (Fig. 1).

**Figure 1.**
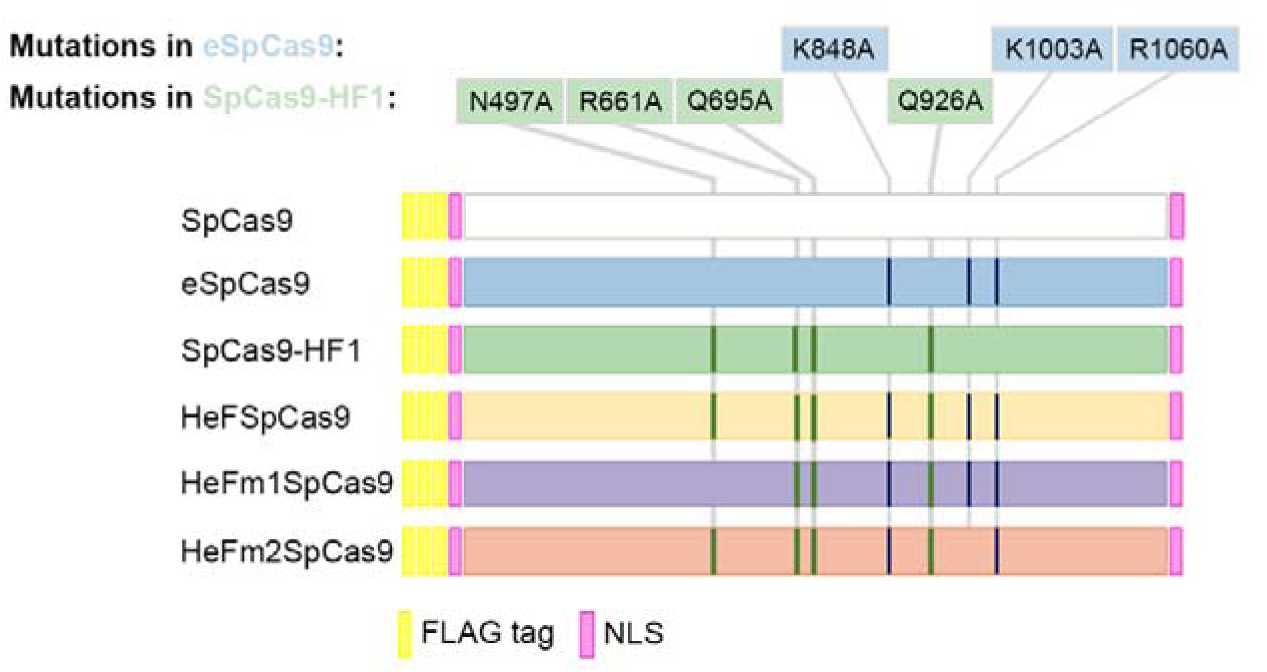
SpCas9 variants employed in these studies. Schematics depicting the main features of the wild type and the five mutant variants of SpCas9 used: each protein sequence is flanked by a nuclear localization signal (NLS) at both ends and is preceded by a 3xFLAG tag. HeF-variants containing combinations of mutations from both eSpCas9 and SpCas9-HF1.

To assess the activity of the Highly enhanced Fidelity SpCas9 nuclease (HeFSpCas9) we performed direct comparisons with eSpCas9, SpCas9-HF1 and the wild type (WT) SpCas9 on a large number of on-target sites. Testing 16 genomic targets in N2a cells, we measured either the on-target HR mediated integration of a GFP cassette with 1,000 bp-long homology arms [35] (Additional File 1: Figure S1a) or the indel frequencies by Tracking of Indels by DEcomposition (TIDE) [36] on five genomic loci (*Pten*, *Prnp*, *Rbl2*, *Ttn* and *Tp53*) (Additional File 1: Figure S1b). To our surprise, the increased fidelity nuclease variants performed poorly in these assays in contrast to those reported earlier [32, 33]. A closer inspection of the results, revealed that many of these targets had been targeted by 21 nt-long spacers bearing a guanine (G) extension at start. This is a commonly applied modification to comply with the preference for a G nucleotide as transcription initiation site for the human U6 promoter [37] that is used for the sgRNAs here and which is the most commonly-used promoter for mammalian sgRNA expression vectors [8].

### 5′-extended sgRNAs diminish the activities of improved fidelity nucleases more with a matching than with a mismatching G nucleotide

The broad applicability of these nucleases depends on how they are able to utilize modified sgRNAs. The need to accept modified sgRNAs is most common because a U6 promoter is used for sgRNA expression, which requires a starting G nucleotide for efficient transcription. According to whether the target requires a G at that position, different practices exist for modifying the 5'-end of the spacer of the sgRNA to meet the starting G requirement. We systematically examined the compatibility of routinely implemented modifications of 5'-ends of the spacers with the SpCas9 variants. In these experiments, we examined 100 sgRNAs using an EGFP disruption assay (Additional File 1: Figure S1c-e) in N2a.EGFP cells [15, 38]. The 20-nt spacers to be examined, unless otherwise noted, start with a 5'-end G nucleotide to facilitate specifically investigating the effect under scrutiny. Fir st we examined the effect of appending either (i) a 21^st^ 5'-end non-matching (Fig. 2a and Additional File 1: Figure S2a and S2c) or (ii) a matching G nucleotide (Fig. 2b and Additional File 1: Figure S2b and S2d) to the spacer sequence on the activities of the increased fidelity nucleases. We compared this activity to that of the wild type and to the corresponding unmodified guides (with 20 nt-long spacer). The results of these experiments (Fig. 2a and b) demonstrate that 21 nt-long spacers interfere with the activities of the increased fidelity nucleases, providing an explanation to the effects previously seen on Additional File 1: Figure S1a and b. Interestingly, extending the guide with a matching 5'-end G nucleotide is much more detrimental to the activities of these nuclease variants than extending it with a mismatching one. In further experiments, we systematically examined and showed, confirming also earlier reports [32, 33], that the routine application of the increased fidelity nucleases is not compatible with modified sgRNAs that are generated by commonly-applied approaches such as (i) altering the non-G 5'-end nucleotide to a G or (ii) using the spacers without alteration, i.e. with the 5'-end non-G nucleotide or (iii) truncating back the guide until a G nucleotide is encountered resulting in spacers of between 19 to 17 nucleotide-length (Additional File 1: Figure S2e and f). All of these alterations diminish the activities of all mutant nucleases, however, to different extents; eSpCas9 showed lower sensitivity compared to SpCas9-HF1. These results indicate that enhanced and high fidelity nucleases are generally not compatible with these approaches and can be used only with matching 20 nucleotide-long spacers. This finding is critical for their effective application and methods that relax the restriction for a 5'-end G nucleotide imposed by employing the U6 promoter for sgRNA expression might prove to be very valuable tools for these nucleases [39-43] (see Additional File 1: Supplementary results 1 for detailed information).

**Figure 2.**
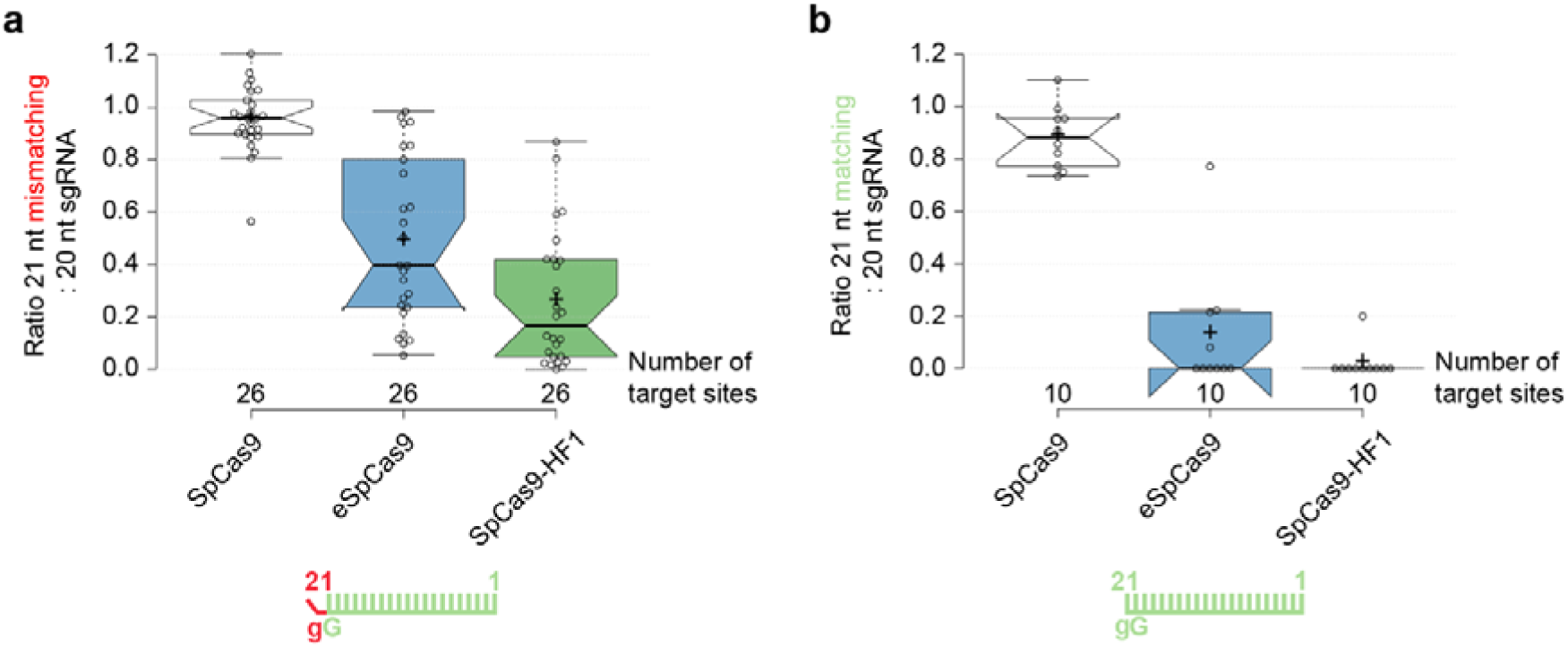
Extending the guide RNA with a matching 5′-end G nucleotide is much more detrimental to the activities than extending with a mismatching one in case of eSpCas9 and SpCas9-HF1. Effect of 5′-extension of the sgRNA with **a**, a mismatching G or **b**, a matching G nucleotide on the activities of SpCas9 nucleases in comparison with using perfectly matching 20 nt-long spacers (data used are from Additional File 1: Figure S2a, c and S2b, d; sites targeted are provided in Additional File 2). Schematics for the spacers used are depicted below the categories as green color combs and the 21^st^ G nucleotide extensions are depicted as a red color bent end teeth if mismatching; lower case g-s represent appended nucleotides; numbering corresponds to the distance from the PAM. Tukey-type notched boxplots by BoxPlotR [67]: center lines show the medians; box limits indicate the 25^th^ and 75^th^ percentiles; whiskers extend 1.5 times the interquartile range from the 25^th^ and 75^th^ percentiles; notches indicate the 95% confidence intervals for medians; crosses represent sample means; data points are plotted as open circles and correspond to the different targets tested (in total 26 and 10, respectively, for a and b). Each pair of means is statistically different at the p<0.05 level: a, SpCas9 – eSpCas9 (<.001), SpCas9 – SpCas9-HF1 (<.001), SpCas9-HF1 – eSpCas9 (<.015); b, Statistically different pairs: SpCas9 – eSpCas9 (<.001), SpCas9 – SpCas9-HF1 (<.001).

### There is a ranking by activity and target discrimination/selectivity among the improved fidelity nuclease variants

Based on the above results, we performed direct comparisons of HeFSpCas9 with eSpCas9, SpCas9-HF1 and the wild type SpCas9, employing 24 sgRNAs with 20 nt-long spacers using EGFP disruption assay. Both eSpCas9 and SpCas9-HF1 demonstrate high activities (Fig. 3a and Additional File 1: Figure S1f), showing >80% activity for more than 83% and 41% of these 24 targets, respectively, when compared to the wild type SpCas9 (Fig. 3b). HeFSpCas9 shows activity only on a subset of these targets. Interestingly, although these nucleases show decreased overall activity, they are capable of undiminished activity, comparable to that of WT SpCas9 on particular individual targets. eSpCas9 is the least discriminative, HeFSpCas9 the most discriminative in target selection in this respect.

**Figure 3.**
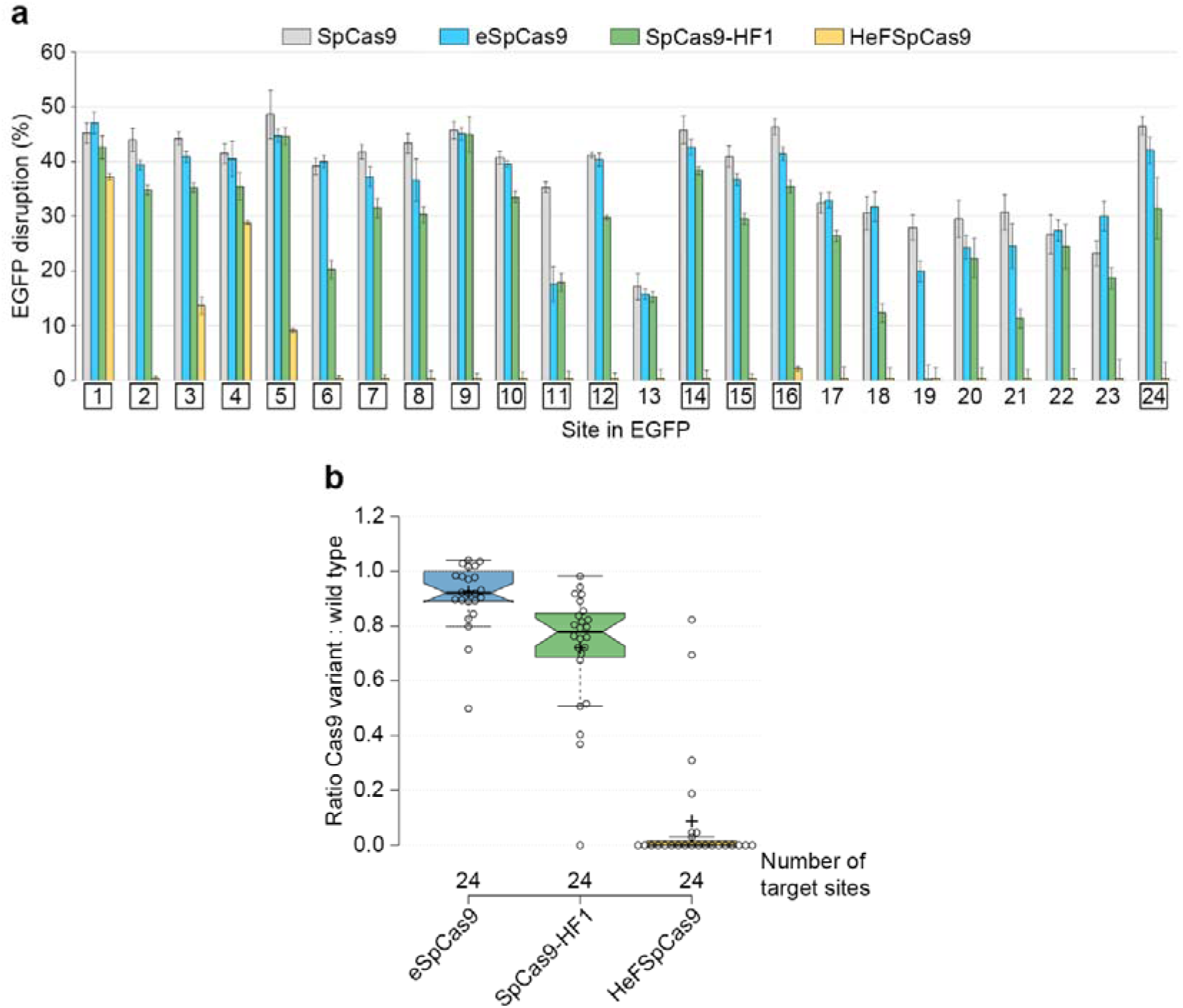
Side-by-side comparison of SpCas9 variants programmed with perfectly matching sgRNAs reveal a target-selectivity ranking among the variants in the order of eSpCas9>SpCas9-HF1>HeFSpCas9. **a**, EGFP disruption activities of the nucleases, calculated as described in Methods. Bars correspond to averages of n=3 parallel samples; error bars represent the standard errors estimated by Gaussian error propagation of the component standard deviations (s.d.-s) associated to both EGFP and mCherry (transfection control) values. The target numbers marked by square are the 16 targets examined on Figure 4a. **b**, Summary of the characteristics of distributions of data for on-target disruption activities of nuclease variants normalized to that of the wild type SpCas9. Tukey-type notched boxplots by BoxPlotR: center lines show the medians; box limits indicate the 25^th^ and 75^th^ percentiles; whiskers extend 1.5 times the interquartile range from the 25^th^ and 75^th^ per centiles; notches indicate the 95% confidence intervals for medians; crosses represent sample means; data points are plotted as open circles. The sample points (24 in case of each variant) correspond to the targets present on Figure 3a. Each pair of means is statistically different at the p<0.05 level: SpCas9-HF1 – eSpCas9 (<.002), SpCas9-HF1 – HeFSpCas9 (<.001), HeFSpCas9 – eSpCas9 (<.001).

### There is a fidelity ranking among the mutant SpCas9 nuclease variants

We also aimed to compare the fidelity of these nucleases. Unbiased genome-wide off-target analyses are reported to barely detect off-targets in case of “typical sequences” [33] cleaved by eSpCas9 and by SpCas9-HF1 [32, 33], making this approach less appropriate to carry through a comparative analysis of the nucleases. Thus, we decided to compare the effect of single base mismatches on their fidelity in an EGFP disruption assay. This assay is a sensitive surrogate for the off-target activity of Cas9 nucleases [15, 38]. We placed mismatches in the PAM distal regions (between positions 19^th^ to 14^th^) of the spacer sequence, where mismatches are most tolerated by SpCas9. Here, 16 targets were examined employing 144 mismatching sgRNAs; each target with a matching and with 9 one-base mismatching sgRNAs (three possible mismatches for each of the three positions, as on Figure 4a). Since we found that mixing the three possible sgRNAs mismatching the same position resulted in sensitive reporting of the off-target activities by the disruption assay (Additional File 1: Figure S3a), we used this approach here.

**Figure 4.**
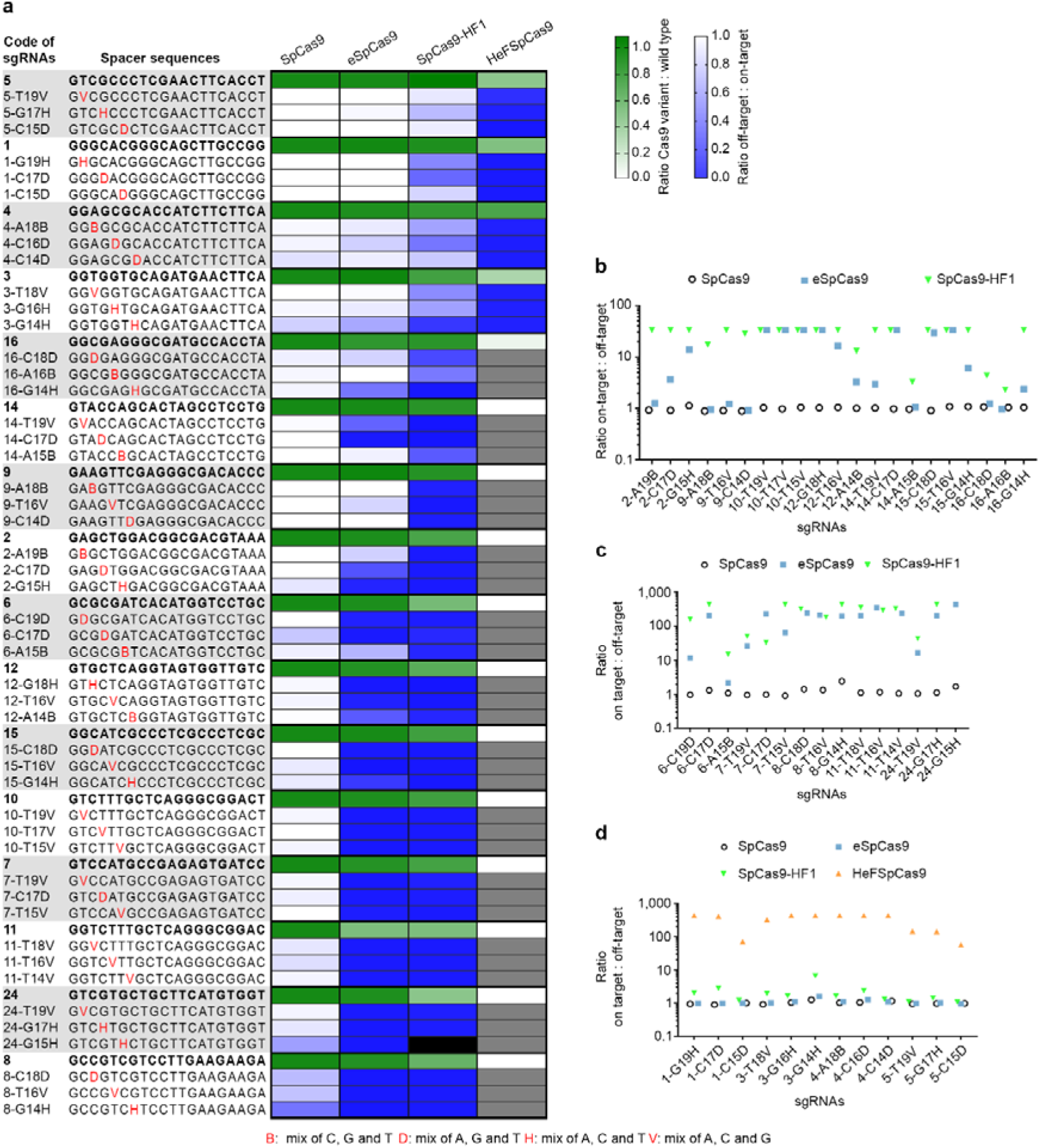
A cleavability-ranking of the targets by the nuclease variants as well as a fidelity ranking (eSpCas9<SpCas9-HF1<HeFSpCas9) of the nucleases on these targets is apparent. Disruption and indel formation activities of SpCas9 nucleases programmed with perfectly matching or partially mismatching sgRNAs. a, Heat maps showing the relative activities (white to green) of the nuclease variants compared to the wild type for each of the targets and the ratios of off-target to on-target disruption activities (blue to white) of the wild type and mutant nucleases measured employing the indicated target and mismatching spacer sequences; grey and black boxes: not determined due to diminished on-target activities and sample loss, respectively. **b, c and d**, Specificities (on-target:off-target ratio) of the nucleases assessed by **b**, disruption activities **c**, deep-sequencing on indel formation (eSpCas9 and SpCas9-HF1) and disruption activities (SpCas9), and **d**, deep-sequencing on indel formation (HeFSpCas9) and disruption activities (SpCas9, eSpCas9, SpCas9-HF1).

These results show that there is a big difference in fidelity between wild type SpCas9 and eSpCas9 or SpCas9-HF1. Whereas wild type SpCas9 barely distinguishes perfect matches from mismatches for the majority of the targets examined, eSpCas9 and SpCas9-HF1 are capable of strong discrimination (Fig. 4 and Additional File 1: Figure S3b). eSpCas9 does not exhibit any detectable cleavage (<3%) with 20 out of 48, whereas SpCas9-HF1 shows no cleavage with 28 out of 47 mismatch positions of sgRNAs. These confirm earlier reports that eSpCas9 and SpCas9-HF1 have greatly increased target fidelity not only when presented with multiple-but also with single-base mismatched sequences. Our results also reveal that SpCas9-HF1 possesses higher fidelity, exhibiting less off-target cleavage than eSpCas9. The number of spacers with mismatching positions that result in higher than 80% of the corresponding disruption levels found with the matching spacers are: 42 for the WT protein, 18 for eSpCas9 and only 3 for SpCas9-HF1 (Additional File 1: Figure S3b). Interestingly, there are targets (target number 1, 3, 4 and 5 on Figure 4a) for which even SpCas9-HF1 shows higher off-target effects, although the difference between the sequence-characteristics of these targets with low and high off-target cleavages is not apparent. Importantly, HeFSpCas9 exhibits high fidelity activity particularly on these targets (Fig. 4a).

Since several off-target positions resulted in disruption levels at around or under the detection limit of the assay, we performed deep sequencing on 67 samples to assess more precisely their respective fidelity on single-base mismatched sequences (for details of the read numbers see: Additional File 2: Deepseq reads). SpCas9-HF1 also proved to be a higher fidelity nuclease than eSpCas9 by deep sequencing, showing considerably lower off-target cleavage (>2-fold) with 7 out of 14 mismatch-position containing guide RNAs (Figure 4c). When examining those targets by deep sequencing (number 1, 3, 4 and 5) for which eSpCas9 and SpCas9-HF1 failed to achieve high fidelity, HeFSpCas9 showed detectable off-target activity only with 4 out of 12 mismatching positions, with the rest exhibiting off-target activity indistinguishable from the background. These results demonstrate a spectacularly increased specificity of HeFSpCas9 on these targets, of about 50-to 400fold over of the two other increased fidelity nucleases (Fig. 4d). Thus, the fidelity ranking of the nucleases is, in order of increasing fidelity, eSpCas9<SpCas9-HF1<HeFSpCas9.

### Ranking among the targets

Applying the four nucleases to a number of targets and exploiting a great number of sgRNAs with mismatching guides (Fig. 4a) revealed another important feature: a ranking among the targets due to differences in the efficiency with which targets are cleaved. Targets efficiently cleaved by a nuclease with both perfectly matching and mismatching guides were likely to be cleaved efficiently by another *higher* fidelity nuclease with perfectly matching sgRNAs but not (or much less) with mismatching ones. In this way targets could be ranked according to their requirements for a type of nuclease to result an optimal (efficient with minimal off-targets), high specificity cleavage. In this ranking, at one end are those targets that are efficiently cleaved by all three nuclease variants, however, only the highest fidelity HeFSpCas9 cleaves them with little to none off-target effect. Such targets are e.g. 1, 3, 4 and 5 in Figure 3 and 4. On this kind of targets both eSpCas9 and SpCas9-HF1 showed significant off-target activities with single-base mismatched guides: in particular they showed on average 93% and 60% of the corresponding on-target activities, respectively, while for the rest of the targets (12/16) these off-target activities were only 26% and 6%, respectively (calculated from the data shown on Figure 4). By contrast, at the other end of the ranks are those targets that are cleaved by eSpCas9 with >80% efficiency of WT SpCas9 (such as targets 8 and 24 on Fig. 4a) and without much off-target activity (note that for such targets even the WT protein has decreased off-target activity). Although the difference in terms of fidelity and target selectivity is much larger between SpCas9-HF1 and HeFSpCas9, target ranking is also clearly discernible between eSpCas9 and SpCas9-HF1, targets 2, 9, 14, 16 being efficiently and much more accurately targeted by SpCas9-HF1.

To confirm that these results are not specific only for the mouse cell line used, we selected 9 targets of various ranks (covering optimal sequences for each of the variants, e-,-HF1 or HeFSpCas9 and used previously in the experiment on Figure 4a) and repeated the same experiment in human cells (HEK-293), by using a cell line containing a single-copy integrated EGFP (HEK-293.EGFP) similarly to the N2a.EGFP cells used. To each target site 9 single-base mismatched guides were applied thus using altogether 81 mismatching guides. The patterns revealed by these experiments on the nucleases (Additional File 1: Figure S4) are almost identical to those found previously (shown on Figure 4a), supporting the idea that the features of the increased fidelity nucleases and that of the targets, which have become apparent in this study, are their intrinsic characteristics and are not specific to the particular cell line used in the experiments. The new observation about target ranking reported here is a key aspect of selecting the optimal variant for a particular target.

Since off-target effects are also dependent on the amount of the active nuclease present, we have also compared the activities as a function of the expression levels of these nucleases while making considerable efforts to match the expression levels of the variants as closely as we could (see Additional File 1: Figure S5, S6 and Supplementary results 2 for more details). These experiments revealed that the relative efficiencies and specificities of these nuclease variants seen in these studies, particularly the differences between eSpCas9 and SpCas9-HF1, are primarily determined by their intrinsic characteristics; however, it cannot be ruled out that there might also be a contribution to the lower on-target activities and higher fidelity of SpCas9-HF1 seen here from its somewhat lower expression levels from these vectors.

From the observed target ranking it should follow that the higher the cleavage activity of a nuclease with one mismatching-position bearing spacer on a target, the higher is its activity with any other type of modified/imperfectly matching (extended, truncated, or with another mismatching position) guide on the same target, if they act using the same mechanism. Such a correlation is discernible from experiments employing sgRNAs with mismatching and extended spacers on the same set of targets (Fig. 4a, Additional File 1: Figure S2a and c). Correlation matrix analysis of the activity data shows positive Pearson correlations (0.83 or 0.78) that are significant between the disruption activities of either eSpCas9 or SpCas9-HF1 programmed with sgRNA-s bearing single mismatching nucleotides and those being extended with a 5'-end mismatching G targeting the same site (Additional File 1: Figure S7a and c). In addition, differing positions of mismatches for the same targets also showed significant positive correlation (between 0.73 to 0.93 and 0.73 to 0.97 for eSpCas9 and SpCas9-HF1, respectively) (Additional File 1: Figure S7b and d). This result further supports the idea of target-ranking revealed but also implies that a similar mechanism determines how these imperfectly matched or modified guides affect the cleavage activities of SpCas9 nucleases.

To confirm the observed nuclease-and target-ranking, an additional 26 spacers were screened for on-target cleavage activity in the EGFP disruption assay. eSpCas9 and SpCas9-HF1 demonstrate decreased activity on some of these targets compared to the WT protein to an extent comparable to that found previously (Additional File 1: Figure S8a and b). Importantly, while HeFSpCas9 shows no detectable activity on 22 out of 26 targets, it shows about 80% of the WT activity on two targets (EGFP site 26 and 43). To test the fidelity of SpCas9-HF1 and HeFSpCas9 when dealing with these two targets, 9-9 sgRNAs (each containing a single-base mismatch in one position) were applied. In line with expectations, SpCas9-HF1 cleaves these targets with non-perfectly matching sgRNAs (Additional File 1: Figure S8c). By contrast, HeFSpCas9 demonstrated considerably less off-target activity in these experiments.

### Generation of a new variant with in-between activity and fidelity

For the majority of the targets examined here one could choose from these high fidelity nucleases one that attains at least 70% disruption activity compared to the wild type protein, combined with a minimal off-target activity on single-base mismatches (Fig. 3, Fig. 4). However, there are a few targets on which even the higher fidelity SpCas9-HF1 showed considerable off-target activity, but were not yet effectively cleaved by HeFSpCas9. Based on the ranking of the nuclease variants and targets established here, we proposed that a variant with on-target and off-target activity in-between that of the two nucleases (HeFSpCas9 and SpCas9-HF1) could be generated by reverting single mutations out of the seven in HeFSpCas9. Based on the results of the former studies [32, 33] we conjectured that an HeFSpCas9 derivative lacking either the N497A mutation of SpCas9-HF1 (HeFm1SpCas9) or the K1003A mutation of eSpCas9 (HeFm2SpCas9) would perform better on these tar gets and gener ated these mutants. We selected 17 targets wher e SpCas9-HF1 showed either off-target cleavage in the disruption assay as shown in Figure 4a (at least with one mismatched-position) or showed higher on-target activity as shown in Additional File 1: Figure S8. We tested these targets with the five nucleases, WT-,-HF1, HeF-, HeFm1-and HeFm2SpCas9. The results obtained suggest that HeFm1SpCas9 is quite similar to HeFSpCas9, whereas HeFm2SpCas9 demonstrated on-target and off-target activities that are more in-between the activities of SpCas9-HF1 and HeFSpCas9: showing higher on-target activities (for 11 out of 17 targets; Fig. 5a and Additional File 1: Figure S9a and b) but also, slightly higher off-target activities (for 2 out of 5 targets; Additional File 1: Figure S9c) than HeFSpCas9. Compared to SpCas9-HF1, HeFm2SpCas9’s specificity is higher (Fig. 5b) for those 5 targets on which it demonstrated more than 60% of the activity of the WT protein as shown in Figure 5a. Thus, we successfully generated a nuclease (HeFm2SpCas9) with fidelity and target-selectivity in-between SpCas9-HF1 and HeFSpCas9. These data suggested that HeFm2SpCas9 might be worth a more thorough characterization and accordingly, we also involved these 2 new mutants (HeFm1-and HeFm2SpCas9) in the subsequent experiment.

**Figure 5.**
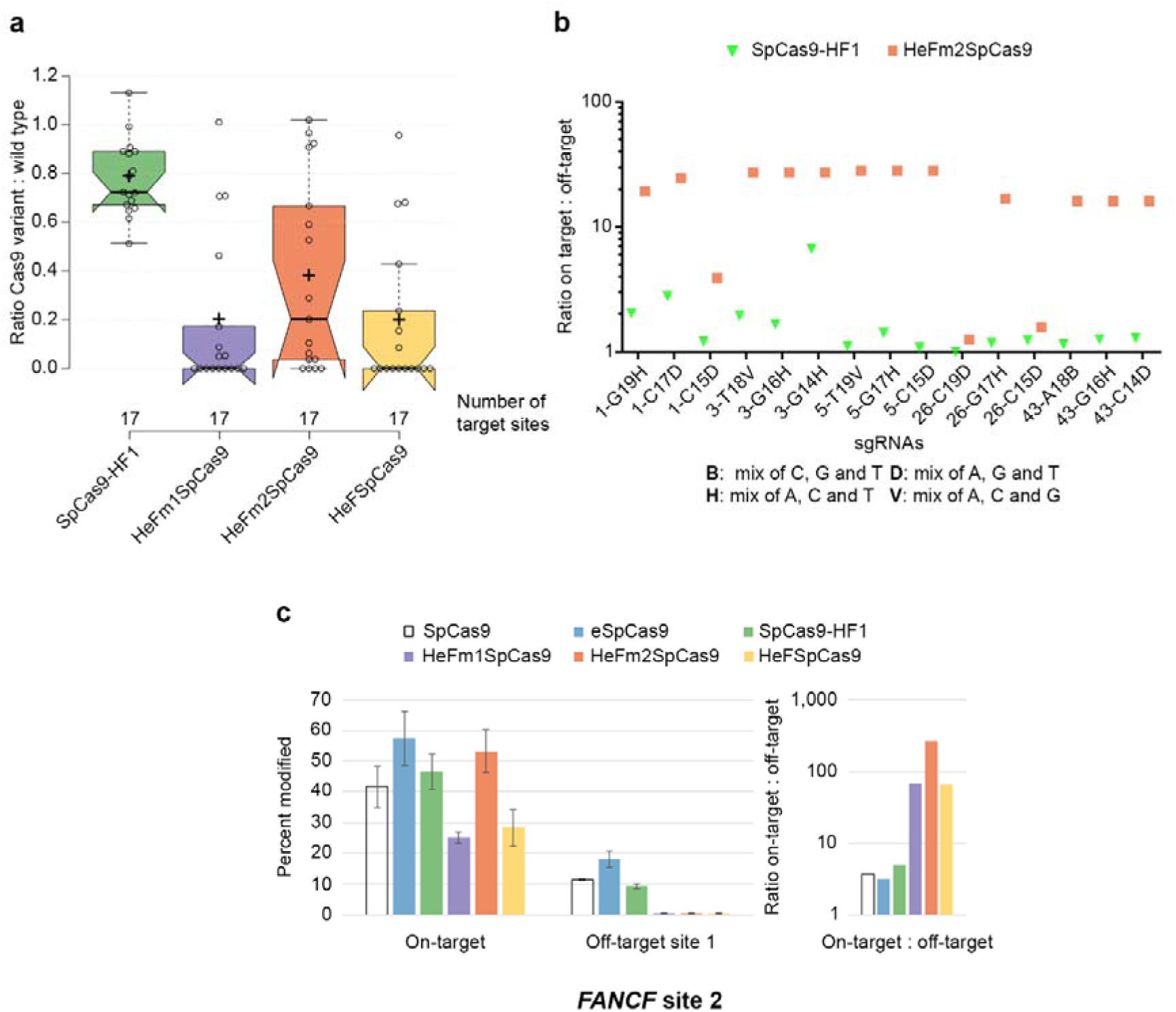
Variant HeFm2SpCas9 exhibits in-between target-selectivity and fidelity of SpCas9-HF1 and HeFSpCas9. Disruption and indel formation activities of SpCas9 nuclease variants bearing combinations of eSpCas9 and SpCas9-HF1 mutations. **a**, Tukey-type notched boxplots of ratios of on-target disruption activities of the variants to those of wild type nucleases as indicated (data used are from Additional File 1: Figure S9a and b; sites targeted are provided in Additional File 2): center lines show the medians; box limits indicate the 25^th^ and 75^th^ percentiles; whiskers extend 1.5 times the interquartile range from the 25^th^ and 75^th^ percentiles; notches indicate the 95% confidence intervals for medians; crosses represent sample means; data points are plotted as open circles (17 in case of each variant) and correspond to the targets tested. Statistically different pairs of means at the p<0.05 level: SpCas9HF1 – HeFm1SpCas9 (<.001), SpCas9HF1 – HeFm2SpCas9 (<.015), SpCas9HF1 – HeFSpCas9 (<.001). **b**, Comparison of specificities of SpCas9-HF1 and HeFm2SpCas9 assessed in disruption assays with partially mismatching sgRNAs on targets where HeFm2SpCas9 reached at least 60% activity of the WT protein on Figure 5a. c, Left panel: mean percent modifications by WT SpCas9 and variants at the FANCF site 2 as well as off-target site 1 from Figure 5 in ref. [33], that is readily cleaved by SpCas9-HF1. Percent modifications are determined by TIDE; error bars represent s.d. with Gaussian error propagation for n=3 parallels. Right panel: Specificity of wild type and mutant variants on the FANCF site 2 plotted as ratio of on-target to off-target activity (calculated from the left panel).

Kleinstiver and coworkers showed that SpCas9-HF1 cleaves efficiently the off-target site 1 of *FANCF* site 2 that bears a mismatch close to the PAM region [33]. We expected that such type of targets are a better match for the higher fidelity HeF nuclease variants and tested the nucleases on this target by measuring on-target and off-target indel generation through TIDE [36]. All HeF-variants showed only background-level cleavage activity at the off-target site, whereas there were differences in on-target activity: among them the activity of HeFm2SpCas9 was the highest and comparable to that of the wild type (Fig. 5c). This supports our conjecture that HeFm2SpCas9 might be a good candidate to fill the gap between SpCas9-HF1 and HeFSpCas9 and demonstrates the relevance of our approach.

### Binding of these nuclease variants to inefficiently-cleaved target DNAs is apparently not diminished as compared to the wild type nuclease

Another application of SpCas9, besides being a programmable nuclease, is using its inactive variant for delivering effector domains precisely to a chosen locus within the genome [28, 44-51]. Even active SpCas9 can be exploited for sequence-specific binding without target cleavage by complexing it with truncated sgRNAs harboring 14-15 nt-long spacer sequences [32, 52, 53]. The study presented on Additional File 1: Figure S2f reveals that increased nucleases exhibit no cleavage activity when employing truncated sgRNAs missing more than two nucleotides. We wondered if these high fidelity nucleases (eSpCas9, SpCas9-HF1 and HeFSpCas9) can also be exploited for transcriptional activation. For this reason, we compared their efficiency using five 14 nucleotide-long spacers with an extra non-matching G in their 5'-end positions, targeting the promoter region of the *Prnp* gene that drives the expression of an EGFP cassette in the N2a.EGFP cell line. Contrary to the above expectations, all four nucleases demonstrated comparable activities, resulting a 15-20-fold activation (Fig. 6a), which is similar to that of the catalytically inactive wild type and mutant variants (dead eSpCas9, dead SpCas9-HF1 and dead HeFSpCas9) with the same sgRNA but with 21 nt-long spacers (4 out of 5 with a mismatching 5' G) (Fig. 6b). These results, although indirect, suggest that the binding of the nucleases to their targets are not impaired with altered sgRNAs.

**Figure 6.**
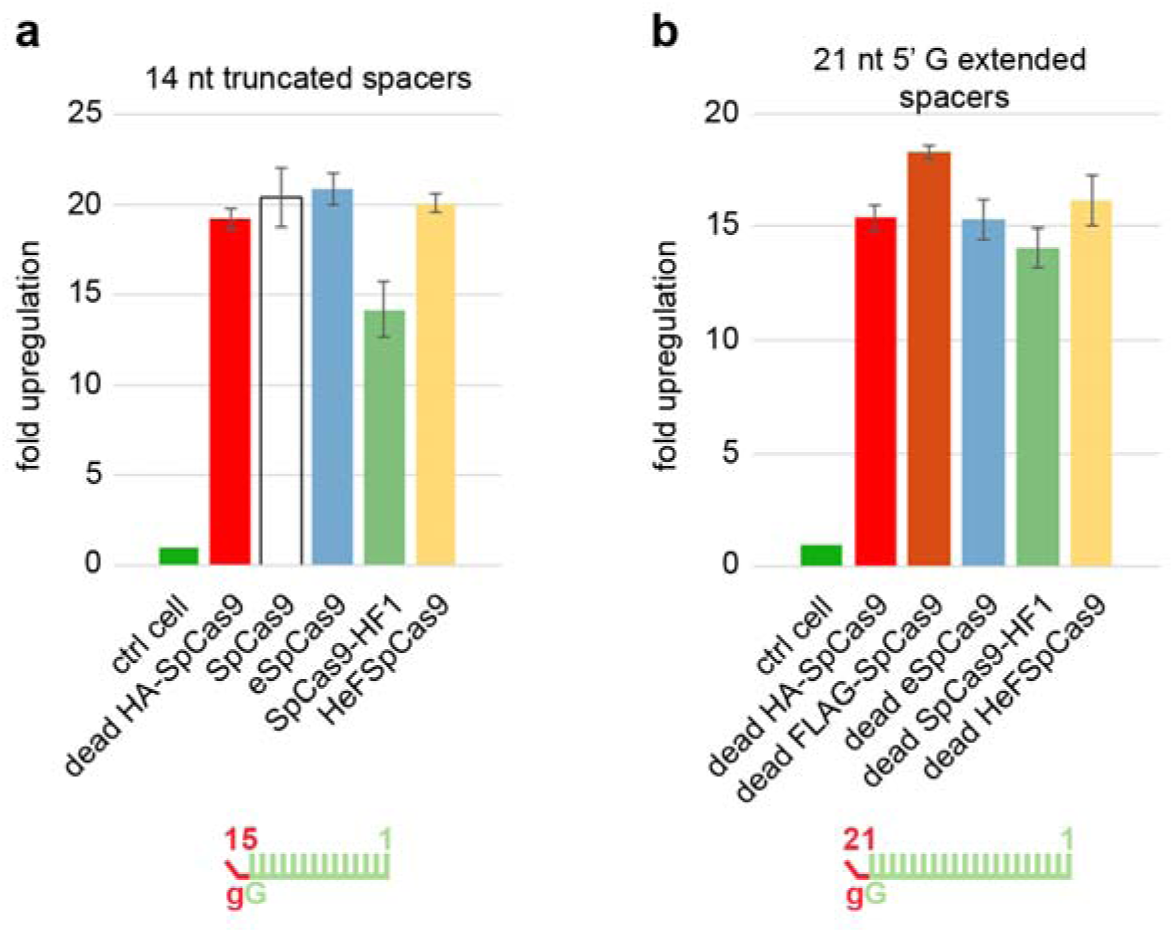
Transcription activation is not impaired compared to the wild type when the nuclease variants are charged with altered sgRNAs. Transcription activation of an EGFP inserted after the *Prnp* promoter, employing five targets in the promoter region. **a**, active nuclease variants programmed with 15 nt-long truncated MS2 aptamer-containing sgRNAs. **b**, Dead nuclease variants programmed with 21 nt-long MS2 aptamer-containing sgRNAs. Spacers used are shown as green combs where green color represents matching, while red color represents mismatching positions; lower case g-s represent appended nucleotides to the 5′-end of the guide; numbering corresponds to the distance from the PAM. Bars correspond to averages of n=3 parallel samples; error bars represent the standard deviations (s.d.).

Employing a more direct in vitro approach we performed polyacrylamide-gel electrophoretic mobility shift assay of the cleaved target-DNAs by the nucleases, exploiting three targets that were shown to be cleaved efficiently by WT SpCas9 but not by HeFSpCas9. By these experiments, we confirmed that although HeF nucleases do not show nuclease activity, they retain most of their DNA-binding abilities to these targets (Additional File 1: Figure S10). Furthermore, to understand the effect of appending an extra G nucleotide to the 5'-end of the guides we examined the in vitro binding of eSpCas9 and SpCas9-HF1 charged with 21 nucleotide-long sgRNAs to selected targets that were only cleaved when the 20 nucleotide-long spacers were applied in the disruption assay (Fig. 2). We found that although the matching G-extension fully diminishes the cleavage activities of these variants on these targets, their binding seems to remain unaffected (Additional File 1: Figure S10).

## Discussion

In principle, there are two types of approaches that have been successfully used to decrease off-target cleavage activities of SpCas9. On the one hand, the length of the target sequence necessary to elicit double strand cleavage may be increased by employing dCas9-FokI [29, 30] or nickases [27, 28]. On the other hand, lessening the promiscuity of SpCas9 can be achieved by lowering its activity on off-target sequences while maintaining reasonably effective on-target activities. This is attempted by minimizing the exposure of the DNA to SpCas9 activity by limiting the time of its expression [24, 25, 54-57] or decreasing the level of protein [58, 59], employing modified sgRNAs (truncated [15] or 5'-extended with a GG dinucleotide [21]) or weakening the protein-DNA interactions [32, 33]. Although a systematic comparison of these approaches is still missing, they have led to varying success and it has been observed that different targets are cleaved with different off-target propensity by SpCas9. Unfortunately, in the absence of sufficiently accurate prediction tool, the off-target propensity of SpCas9 on a given target can be determined only by time-consuming and laborious genome-wide off-target analysis, apart from when homopolymeric and repetitive sequences are encountered, which have been already reported to be cleaved with high off-target effects, and hence could be easily avoided. For the targeting of sequences in the bulk of the genome our results take a step forward in overcoming these difficulties.

The most important outcomes of the experiments reported here are (i) the observed fidelity or activity and target selectivity ranking among the increased fidelity nucleases and (ii) a ranking of the cleavability of targets without off-target effects by these nuclease variants. These results may be understandable now; however, before this study it was not clear [32, 33] whether the mutations in eSpCas9 or SpCas9-HF1, although aiming at non-specific contacts, altered their sequence specificity, i.e. if eSpCas9 and SpCas9-HF1 exhibit lower activities on distinct sets of targets, a result which would not support the existence of the target-ranking discerned here. It was also not clear whether SpCas9-HF1 had higher fidelity than eSpCas9 and whether off-target propensities varied according to the target.

According to our findings, eSpCas9 possesses the lowest target selectivity among the increased fidelity nucleases examined here. It has activity comparable to that of WT SpCas9 when employed with 20 nt-long, perfectly matching sgRNAs and because of its higher fidelity its routine use is preferable to using WT SpCas9, for practically all applications. We did not find a single target for which WT SpCas9 shows higher specificity (defined as on-target/off-target ratio) than eSpCas9. In contrast, SpCas9-HF1 showed strongly decreased activities for some targets (such as targets 6, 18, 19 and 21 on Fig. 3a and 27, 40, 45 and 50 on Additional File 1: Figure S8a) in these experiments. Even though it exhibits markedly higher specificity for several targets (e.g.: targets 2, 9, 14 and 16 on Fig. 4) than eSpCas9, it may not be advisable to use it routinely for any targets without pretesting. Kleinstiver et al. suggested that SpCas9-HF1 cleaves only “atypical”, repetitive or homopolymeric targets with substantial off-target propensity [33]. According to our experiments, some typical targets that are not homopolymeric or repetitive sequences are also cleaved with higher off target propensity by SpCas9-HF1. The proportion of such targets exceeds ten percent occurrence in our experiments. Unfortunately, at present we are not able to predict which targets would have a higher off-target propensity, which limits the unrestricted use of SpCas9-HF1 for DNA modifications aimed at avoiding off-target effects. The HeF nucleases developed here exhibit greatly increased fidelity and an accompanying increased target-selectivity/lower activity; a finding that may not be surprising. However, we found it striking that they exhibit this high fidelity specifically for those targets for which SpCas9-HF1 fails to show improved fidelity. This suggests that HeF-variants will be very useful complements to the already existing increased fidelity nucleases in genome engineering applications, perfectly fulfilling the anticipated role for which we generated them.

Another interesting result is the finding that 5'-extending the sgRNAs with a single G nucleotide diminishes the activities of the increased fidelity SpCas9 variants. A 5' GG dinucleotide extension of the sgRNA has been reported to decrease the off-target cleavage propensity of SpCas9 with certain targets [21]. Since the 5'-end of the sgRNAs protrudes relatively freely from the known SpCas9-sgRNA structures [9](Additional File 1: Figure S11a and b), the mechanism by which the 5' dinucleotide extension decreases off-target propensity was not apparent and it was suggested that it may alter the guide RNA’s stability, concentration, or secondary structure [21]. More recently a structure of SpCas9 has been published, which provides the most detailed picture of a cleavage competent state of the SpCas9-sgRNA complex [60]. According to this structure the 5'-end of the sgRNA is not only buried inside the protein, but if extended it could only exit the protein’s structure at a separate opening that is at some distance from the target DNA strand and separated from it by parts of the protein (Additional File 1: Figure S11c, d and e). Such a structural arrangement would explain how a 5' G extension may disturb the cleavage but not the binding of SpCas9. It also makes it understandable how a matching G extension that would lengthen the RNA:DNA hybrid helix may diminish much more the cleavage activities of SpCas9 than a mismatching G may, by causing larger distortions to the cleavage-competent structure.

The results of these studies also extend our knowledge of the factors that influence on-target and off-target cleavages of SpCas9 and its increased fidelity variants in several respects. High fidelity nucleases exhibit strongly reduced activities with truncated sgRNAs leading to the conclusion that they are likely to improve specificity by a similar mechanism to that of decreasing the interaction energy of the protein–sgRNA complex or R-loop stability [22, 23]. Our results may further complement this picture. (i) Increased fidelity nuclease variants show decreased activity with 5'-extended guides as well, suggesting a similar mechanism of action between extending (or truncating) the guide, and variants engineered to have reduced non-sequence specific DNA contacts. (ii) This contention of similar mechanisms is further strengthened by the correlations we found between the activity-reducing effect of appending a 5' G to the sgRNAs and of using imperfectly matching sgRNAs for cleaving the same target sequences with an increased fidelity nuclease (Additional File 1: Figure S7). (iii) These correlations also show that highly reduced off-target effects vary according to the target, an effect that we term “off-targetless cleavability ranking” and is also apparent in Figure 4a. (iv) Furthermore, we show here that increased fidelity nucleases using sgRNAs that are incapable of cleaving the DNA are able to elicit transcription activation and bind DNA in vitro similarly to the WT protein (Fig. 6 and Additional File 1: Figure S10). Altogether these observations suggest that these modifications, truncating or extending the guide and reducing non-sequence specific protein-DNA contacts improve SpCas9 fidelity by primarily affecting the catalytic activity of the cleavage complex without much altering the target binding of SpCas9.

The most simple interpretation of these data is one similar to that originally proposed to explain TALEN off-target activity [23, 61, 62]. However, in case of SpCas9, the specificity of its cleavage activity is determined by the interaction energy of its cleavage complex rather than by its binding. In this scenario, the actual sequence of a target also contributes to the energy of the cleavage complex. With high energy-contributing targets, the cleavage complex may possess enough excess energy to tolerate mismatches in the target DNA:RNA hybrid helix. With low-energy contributing targets the cleavage complex has no or has less excess energy, thus, it is less prone to off-target cleavage. Approaches such as employing various increased fidelity nuclease variants, truncating or extending the sgRNAs decrease the energy of the cleavage complex to different extents: in some cases retaining considerable off-target propensity, or in other cases abolishing even the on-target activity, whereas in optimal cases eliminating most or all of the off-target-, while retaining high on-target cleavage activity.

Consequently, our data point to the fruitlessness of attempting to generate a “generally superior” SpCas9 nuclease variant with overall highest specificity. Rather, an optimal nuclease variant needs to be identified for each target; this knowledge is critical for achieving further minimization of unwanted, off-target cleavage of the genome. Our data suggest that the fidelity of the improved nuclease variants might be further increased by combining them with a nickase approach. However, since the binding activity of these increased fidelity nucleases does not seem to decrease for targets not being cleaved by the given variant (Additional File 1: Figure S10), a combination with a dSpCas9-FokI approach is less likely to be rewarding.

Our results on the use of 5' G-extended sgRNAs (Fig. 2) also offer an alternative solution for increasing the fidelity of WT SpCas9 cleavage. Given the fact that extending the sgRNA with a G nucleotide only modestly decreases the activity of WT SpCas9, (10% on average [Fig. 2]), on the basis of the interpretations provided here, we predict a considerably decreased off-target cleavage with such targets. Furthermore, it is plausible to speculate that in contrast to the so called tru-sgRNAs [15], a matching G-extension may not generate new off-target sites that are not detected using the same unmodified sgRNA-WT SpCas9 complex. In addition, it is likely to increase the fidelity of WT SpCas9 more than non-matching 5' G extensions; two ideas which need further confirmation.

One of the biggest challenges on the field remains to develop methods for the prediction of the ranking of the targets and their matching to the optimal off-target minimizing strategy. Both the approaches we developed here and unbiased genome-wide off-target analyses are excessively laborious and time-consuming, and thus, it is not feasible to apply them to reveal the targets’ cleavability ranking for routine use of SpCas9. However, the ranking of the increased fidelity nucleases observed here, offers a straightforward solution to this problem: a simple pretesting of the candidate target with eSpCas9, SpCas9-HF1 and HeFm2SpCas9 would reveal which is the highest fidelity nuclease exhibiting sufficient activity for the given application.

For the therapeutic usage of the SpCas9-based technology it is of paramount priority to reduce the possibilities for incidental off-targets to a minimum, even below the detection limit of the currently existing approaches (<0.1% imposed by the current NGS technology). These applications frequently involve the optimization of a procedure based on exploiting one or a few targets that are at appropriate positions for their later routine use. The rankings observed here among these nucleases and among targets, and the extremely high fidelity of these increased fidelity nucleases on certain targets suggest that by careful matching of the nuclease to the target being exploited, the off-target cleavage may be reduced even further to the level of incidentally-occurring off-targets that are not detectable by the current methods. These, in turn, would greatly reduce the whole genome sequencing effort required to validate engineered cells for clinical applications without unwanted genomic alterations.

## Conclusion

Increased fidelity nucleases can routinely only be used with perfectly matching 20 nucleotide-long spacers. A 5' G extension of the sgRNAs is especially detrimental for increased fidelity nucleases if the appended G matches the target diminishing their cleavage activity. A fidelity and target selectivity ranking of the increased fidelity SpCas9 nucleases and a cleavability ranking of targets could be revealed here due to our special experimental design: application of SpCas9 variants with significantly different fidelities compared side by side on the same set of targets using a number of imperfectly matching sgRNAs. These experiments revealed that eSpCas9 has lower fidelity than SpCas9-HF1; however, owing to its low target-selectivity it may be used instead of WT SpCas9 for higher fidelity gene editing in most of applications. Interestingly, contrary to expectations, the higher fidelity SpCas9-HF1 also exhibited considerable off-target propensity not only on "atypical" but on "typical" targets that feature no homopolymeric or repetitive sequences. By combining the mutations of these two increased fidelity nucleases, we generated HeFSpCas9, a highly enhanced fidelity nuclease, that is a very useful complement to SpCas9-HF1 by cleaving exactly those targets with little to none off-target effects (with a considerable on-target activity) for which SpCas9-HF1 presented lower fidelity. The observed ranking of the nucleases offers straightforward means for finding the optimal nuclease variant for efficient on-target and minimal off-target modifications. Our approach also provides a framework for the generation of new high fidelity nuclease variants with fidelity in between that of the existing ones, as it is demonstrated here by the development of HeFm2SpCas9. This approach allows "optimal" increased fidelity nuclease variants to tailored to individual targets, matching them to achieve accurate genome editing with minimal off-target effects.

## Methods

### Materials

Restriction enzymes, Klenow polymerase, T4 ligase, Dulbecco’s modified Eagle Medium DMEM (Gibco), fetal bovine serum (Gibco), Turbofect, TranscriptAid T7 High Yield Transcription Kit and penicillin/streptomycin were purchased from Thermo Fischer Scientific, protease inhibitor cocktail was purchased from Roche Diagnostics. DNA oligonucleotides and GenElute HP Plasmid Miniprep kit used in plasmid purifications were acquired from Sigma-Aldrich. Q5 High-Fidelity DNA Polymerase was from New England Biolabs Inc. NucleoSpin Gel and PCR Clean-up kit was purchased from Macherey-Nagel. All plasmid constructs and PCR products were sequenced by Microsynth AG.

### Plasmid construction

Vectors were constructed using standard molecular biology techniques including one-pot cloning method [63], E. coli DH5α-mediated DNA assembly method [64] and Body Double cloning method [65]. For detailed cloning and sequence information see Additional File 3. sgRNA target sites and mismatching sgRNAs are available in Additional File 2. The sequences of all plasmid constructs were confirmed by Sanger sequencing.

Plasmids acquired from the non-profit plasmid distribution service Addgene (http://www.addgene.org/) are the following: pX330-U6-Chimeric_BB-CBh-hSpCas9 (Addgene #42230) [8], eSpCas9(1.1) (Addgene # 71814) [32], VP12 (Addgene #72247) [33], pX335-U6-Chimeric_BB-CBh-hSpCas9n(D10A) (Addgene #42335) [8], pET-dCas9-VP64-6xHis (#62935) [58], sgRNA(MS2) cloning backbone (Plasmid #61424) [51], MS2-P65-HSF1_GFP (Plasmid #61423) [51].

Plasmids developed by us and are deposited at Addgene are the following:

pX330-Flag-wtSpCas9 (without sgRNA) (Addgene #92353), pX330-Flag-eSpCas9 (without sgRNA) (Addgene #92354), pX330-Flag-SpCas9-HF1 (without sgRNA) (Addgene #92102), HeFSpCas9 (Addgene #92355), HeFm1SpCas9 (Addgene #92110), HeFm2SpCas9 (Addgene #92111) pX330-HA-dSpCas9 (Addgene #92112), pX330-Flag-dSpCas9 (Addgene #92113), pX330-Flag-deSpCas9 (Addgene #92114), pX330-Flag-dSpCas9-HF1 (Addgene #92115), pX330-Flag-dHeFSpCas9 (Addgene #92116) pET-deSpCas9-VP64-6xHis (Addgene #92117), pET-dSpCas9-HF1-VP64-6xHis (Addgene #92118), pET-dHeFSpCas9-VP64-6xHis (Addgene #92119) pmCherry_gRNA (Addgene: #80457) U6-sgRNA(MS2)_EF1a-MS2-P65-HSF1 (Addgene #92120)

### Cell culture and transfection

N2a (neuro-2a mouse neuroblastoma cells, ATCC – CCL-131) cells, N2a.EGFP and HEK-293.EGFP cells (both cell lines containing a single integrated copy of an EGFP cassette driven by the Prnp promoter) and HeLa (ATCC – CCL-2) cells were grown at 37 °C in a humidified atmosphere of 5% CO_2_ in high glucose Dulbecco’s Modified Eagle medium (DMEM) supplemented with 10% heat inactivated fetal bovine serum, 4 mM L-glutamine (Gibco), 100 units/ml penicillin and 100 μg/ml streptomycin. N2a, N2a.EGFP and HEK-293.EGFP cells were plated one day prior to transfection on 48-well plates at a density of approximately 30,000 cells (35,000 cells in the case of HEK-293.EGFP). HeLa cells were plated one day prior to transfection in 24-well plates at a density of approximately 60,000 cells. Transfections were performed with TurboFect transfection reagent according to the manufacturer’s recommended protocol and as detailed below for each corresponding assay performed. Transfections were performed in triplicates unless otherwise noted.

### Flow cytometry

Flow cytometry analyses were carried out on Attune Acoustic Focusing Cytometer (Applied Biosystems) or on CytoFLEX Flow Cytometer (Beckman Coulter). For data analysis Attune Cytometric Software v.2.1.0 and CytExpert 2.0 were used. Viable single cells were gated based on side and forward light-scatter parameters and a total of 5,000-10,000 viable single cell events were acquired in all experiments. Attune Acoustic Focusing Cytometer parameters: the GFP fluorescence signal was detected using the 488 nm diode laser for excitation and the 530/30 nm filter for emission, the mCherry fluorescent signal was detected using the 488 nm diode laser for excitation and a 640LP filter for emission. CytoFLEX Flow Cytometer parameters: the GFP fluorescence signal was detected using the 488 nm diode laser for excitation and the 525/40 nm filter for emission, the mCherry fluorescent signal was detected using the 638 nm diode laser for excitation and a 660/20 filter for emission.

### EGFP disruption assay

N2a.EGFP and HEK-293.EGFP cells were co-transfected with two types of plasmids: SpCas9 expression plasmid (137 ng) and sgRNA and mCherry coding plasmid (78-97 ng) using 1 μl TurboFect reagent per well in 48-well plates. Where indicated, less SpCas9 plasmid variants were used and their amounts are completed to 137 ng by a mock plasmid with identical size. Transfected cells were analysed ~72 h and ~168 h post-transfection by flow cytometry. Transfection efficacy was calculated via mCherry expressing cells measured ~72 h post-transfection. EGFP positive cells were counted both ~72 h and ~168 h post-transfection. Background level of EGFP for each experiment was determined on non-transfected cells and also on two types of control transfected population, (i), using co-transfection of a mock plasmid (pmCherry_sgRNA) and an active SpCas9 plasmid or (ii), using co-transfection of a dead SpCas9 expression plasmid and a targeted sgRNA and mCherry coding plasmid.

Percentage of EGFP disruption in the case of N2a.EGFP cells was calculated as follows. The measured percentage of EGFP positive cells for each sample was subtracted from the total average obtained on the controls and was weighted by its transfection efficiency factor. The transfection efficiency factor was obtained utilizing the mCherry present in the samples: by measuring the percentage of mCherry positive cells in individual samples and calculating their deviation from the average percentage of mCherry positive cells obtained on all the transfected samples/wells as: (aver age mCher r y – sample mCher r y)/(aver age mCher r y). For each sample thr ee par allels wer e processed and their values were averaged. The errors associated to the final values were estimated by taking into account the errors of each term, mCherry and EGFP controls as well, using Gaussian error propagation on the standard deviations. When data were further used for ratio plotting, the errors were also processed further by Gaussian propagation to yield the final represented error bars. In the case of HEK-293.EGFP cells the mCherry fluorescence was not used to normalize the data.

### HR mediated integration assay

Cells were co-transfected with three types of plasmids: an expression plasmid for EGFP flanked by 1,000 bp-long homology arms to the *Prnp* gene (referred to as *Prnp*.HA-EGFP plasmid; Tálas et al.[66]) (166 ng), SpCas9 expressing plasmid (42 ng) and an sgRNA/mCherry coding plasmid (42 ng), giving 250 ng total plasmid DNA, using 1 μl TurboFect reagent per well on 48 well plates.

Transfected cells were analysed ~72 h and 12 days post-transfection by flow cytometry. Transfection efficacy was calculated via mCherry expressing cells measured ~72 h post-transfection. EGFP positive cells were counted 12 days post-transfection on three parallels. Background level of EGFP was determined by control co-transfection using dead SpCas9 expression plasmid, *Prnp*.HA-GFP plasmid and a mouse *Prnp* site 1B targeting-sgRNA/mCherry coding plasmid. Values obtained for each sample well were weighted to count for their transfection efficiency utilizing the sample’s mCherry fluorescence and the average mCherry fluorescence value obtained based on all transfected wells as described in EGFP disruption assay. The weighted values of the samples were averaged for parallels and wer e cor r ected for the contr l aver age values. The er r or was estimated by Gaussian er r or propagation of the errors (s.d.) associated with each term experimentally determined that was used for calculation of the value.

### TIDE

Tracking of Indels by DEcomposition (TIDE) method [36] was applied for analysing mutations and determining their frequency in a cell population using different sgRNAs and SpCas9 proteins.

N2a cells were co-transfected with 137 ng of SpCas9 expressing plasmid and 97 ng of sgRNA and mCherry coding plasmid (250 ng total plasmid DNA) using 1 μl TurboFect reagent per well on 48 well plates.

HeLa cells were co-transfected with 400 ng of SpCas9 expressing plasmid and 600 ng of sgRNA/ mCherry coding plasmid (1 μg total plasmid DNA) using 2 μl TurboFect reagent per well on 24 well plates.

Control samples were made for each different genomic target site by co-transfecting a dead SpCas9 expression plasmid and the targeted sgRNA and mCherry coding plasmid.

Transfected cells were divided ~72 h post-transfection as follows: 20% of the cells were analysed for transfection efficacy via mCherry fluorescence by flow cytometry and from the rest of the cells genomic DNA was extracted by following the Gentra DNA Purification protocol (Gentra Puregen Handbook, Qiagen) from the mix of the triplicates. From the isolated genomic DNA PCR was conducted with Q5 High-Fidelity DNA Polymerase (for PCR primer details, see Additional File 2). Genomic PCR products were gel excised (in case of experiments on Additional File 1: Figure S1b) or directly purified (in case of experiments on Fig. 5c) via NucleoSpin Gel and PCR Clean-up kit and were Sanger sequenced. Indel efficiencies were analysed by TIDE webtool (https://tide.nki.nl/) by comparing Cas9 treated and control samples.

### Indel analysis by next-generation sequencing (NGS)

Off-targets with low cleavage efficacy in the EGFP disruption assay together with their on-targets were examined via targeted resequencing. The genomic DNA was extracted at ~ 7 days post-transfection from the mix of the triplicates by following the Gentra DNA Purification protocol. PCR fragments for NGS analysis were generated in two step PCR reactions. Briefly, first step PCR primers (Additional File 2) contained both the *PCR handles* for the second-round amplification and the target specific sequence to amplify genomic regions of interest. The second 10 cycle PCR step, the purification (Ampure Bead clean up), the pooling, the gel excised purification and the 150-bp single-end sequencing on an Illumina MiSeq instrument were performed by Microsynth AG.

### Western blot

N2a.EGFP cells were cultured on 48-well plate and transfected as described above in EGFP disruption assay section. Three days post-transfection, 8 parallel samples corresponding to each type of Cas9 transfected were washed with PBS, then trypsinized and mixed, and were analysed for transfection efficacy via mCherry fluorescence level by using flow cytometry. The cells from the mixtures were pelleted (at 200 rcf for 5 min at 4 °C). Pellet was resuspended in ice cold Harlow buffer (50 mM Hepes pH 7.5; 0.2 mM EDTA; 10 mM NaF; 0.5% NP40; 250 mM NaCl; Proteinase Inhibitor Cocktail 1:100; Calpain inhibitor 1:100; 1 mM DTT) and lysed for 20-30 min. The cell lysates were centrifuged at 19,000 rcf for 10 min. The supernatants were transferred into new tubes and total protein concentration was measured by Bradford protein assay. Before SDS gel loading, samples were boiled in Protein Loading Dye for 10 min at 95 °C. Proteins were separated by SDS-PAGE using 7.5% polyacrylamide gels and were transferred to PVDF membrane, using a wet blotting system (Bio-Rad). Membranes were blocked by 5% non-fat milk in Tris buffered saline with Tween20 (TBST) (blocking buffer) for 2 h. Blots were incubated overnight at 4 °C with primary antibodies [anti-FLAG (F1804, Sigma) at 1:1,000 dilution; anti-β-actin (A1978, Sigma) at 1:4,000 dilution in blocking buffer]. The next day after washing steps in TBST the membr anes wer e incubated for 1 h with HRP-conjugated secondary anti-mouse antibody 1:20,000 (715-035-151, Jackson ImmunoResearch) in blocking buffer. The signal from detected proteins was visualized by ECL (Pierce ECL Western Blotting Substrate, Thermo Scientific) by CCD camera (Bio-Rad ChemiDoc MP).

### Transcriptional activation

N2a.EGFP cells were co-transfected with three types of plasmids as follows: 91 ng of SpCas9 expressing plasmid, 83 ng of the mixture of 5 sgRNA coding plasmid, which expresses MS2-p65-HFS1 fusion protein as well and 75 ng of mCherry coding plasmid (pcDNA3-mCherry) using 1 μl TurboFect reagent per well in 48-well plates. Transfected cells were analysed ~72 h post-transfection. The transfection efficacy was calculated via mCherry fluorescence level. The relative upregulation was calculated from the median of the EGFP intensity. Background level of EGFP was determined using a negative control for transfections: a mock plasmid ([MS2-p65-HFS1_sgRNA(MS2)] without spacer sequence) co-transfected with SpCas9 coding plasmid and with mCherry coding plasmid. The medians obtained for EGFP fluorescence were averaged between three parallel samples and the error was estimated by Gaussian error propagation of the component errors (s.d.) associated to the measured variables.

### In vitro transcription

sgRNAs were in vitro transcribed using TranscriptAid T7 High Yield Transcription Kit and PCR-generated double-stranded DNA templates carrying a T7 promoter sequence. Primers used for the preparation of the DNA templates are listed in Additional File 2. sgRNAs were quality checked using 10% denaturing polyacrylamide gels and ethidium bromide staining.

### Protein purification

pET-dCas9-VP64-6xHis (#62935) was acquired from Addgene, the other Cas9 variants (pET-deSpCas9-VP64-6xHis, pET-dSpCas9-HF1-VP64-6xHis, pET-dHeFSpCas9-VP64-6xHis) were subcloned by us (for detailed cloning information and sequence information see Plasmid construction section and Additional File 3). The resulting fusion constructs contained an N-terminal hexahistidine (His6) tag.

The expression constructs of the dead Cas9 variants were transformed into BL21 Rosetta 2 (DE3) cells and were processed similarly, as follows. Cells were grown in LB medium at 37 °C for 5 h. 10 ml from this culture was inoculated into 1 l of Terrific Broth growth media and cells were grown at 37 °C to a final cell density of 0.6 OD600, and then were chilled to 18 °C. The protein was expressed at 18 °C for 16 h following induction with 0.2 mM IPTG. The protein was purified by a combination of chromatographic steps by using an NGC Scout Medium-Pressure Chromatography Systems (Bio-Rad). Briefly, the cells were harvested and resuspended in 30 ml of Lysis Buffer (40 mM Tris pH 8.0, 500 mM NaCl, 20 mM imidazole, 1 mM TCEP) supplemented with protease inhibitor cocktail (complete, EDTA-free) and were sonicated on ice. The lysate was cleared by centrifugation at 18,000 rcf for 40 min at 4 °C. The supernatant was bound in batch to a 5 ml Mini Nuvia IMAC Ni-Charged column (Bio-Rad). The resin was washed extensively with 40 mM Tris pH 8.0, 500 mM NaCl, 20 mM imidazole, and the bound protein was eluted by 40 mM Tris pH 8.0, 250 mM imidazole, 150 mM NaCl elution buffer. The protein was dialyzed 2x1 h against 20 mM HEPES pH 7.5, 150 mM KCl, 1 mM DTT, 1% glycerol. The dialyzed protein was purified on a 3x1 ml Bio-Scale Mini Macro-Prep High S column (Bio-Rad), eluting with 1 M KCl, 20 mM HEPES pH 7.5, 1 mM DTT. The protein was further purified by size exclusion chromatography on a Superdex 200 16/60 column in 20 mM HEPES pH 7.5, 150 mM KCl and 1 mM DTT. The eluted protein was tested on SDS-PAGE and Coomassie brilliant blue r-250 staining and was stored at −20 °C until use.

### Electrophoretic mobility shift assay

Binding assays were performed in buffer containing 20 mM HEPES pH 7.5, 100 mM KCl, 5 mM MgCl_2_, 0.1 mg/ml heparin, and 1 mM TCEP in a total volume of 20 μl. The sgRNA was supplied at two times the molar amount of protein. The target DNA (40 nM) was incubated with protein-sgRNA complex (160 nM) (or in case of the serial dilution experiments with 160, 320, 640 and 1280 nM protein-sgRNA complex concentrations, respectively) for 30 min at 37 °C. Samples were resolved at 4 °C on an 8% native polyacrylamide gel containing 0.5X TBE and 10 mM MgCl_2_. The gel was stained with ethidium bromide. (For detailed information about sgRNAs and DNA targets see Additional File 2.)

### Statistics

Differences between samples were tested by using Welch’s one-way Anova with Games-Howell post hoc tests for samples with unequal variances or/and samples size (Figures: Fig. 2a; Fig. 2b; Fig. 3b; Fig. 5a; Additional File 1: Figure S2b; S3b; S3b; S8b) and by one-way Anova with Tukey’s post-hoc test for homoscedastic samples (Figures: Fig. 3b; Additional File 1: Figure S1f; S2a; S2e). Homogeneity of variances was tested by Levene’s test. Statistical tests were performed using R version 3.4.1 (C: 2017 The R Foundation for Statistical Computing) and packages FSA, Car, multcomp, userfriendlyscience, dplyr, ggplot2. Test results are in Additional file 4.

The investigators were not blinded to group assignment and outcome assessment.

## Declarations

### Availability of data and material

Plasmids used in this study are available at Addgene: details in Plasmid Construction section. Sequences of the constructs are listed in Additional File 3. The deep sequencing data from this study have been submitted to the NCBI Sequence Read Archive (SRA; http://www.ncbi.nlm.nih.gov/sra/) under accession number: SRR5005491.

### Funding

This work was supported by the National Research, Development and Innovation Office (KFI_16-1-2016-0240).

### Ethics approval and consent to participate

Ethics approval was not needed for the study.

## Acknowledgement

We thank Bernadett Czene, Ildikó Szűcsné Pulinka, Luca Pájer-Turgyán, Anita Paulovitsné Csatári, Dávid Fetter for their excellent laboratory assistance, Lőrinc Pongor, Adrienn Éva Borsy, Márk Saskőy, Éva Varga, Dániel Bátora, István Vida, György Várady, Tamás Hegedűs, Antal Nyeste for their valuable help.

### Author contributions

P.I.K. and E.W. conceived of and designed experiments, P.I.K., A.T., E. T., K.H., Z. L. and E. F. performed all experiments. P.I.K., A.T., E. T., K.H., Z. L. and E. F. analysed the data. P.I.K., E. F., and E.W. wrote the manuscript with input from all the authors.

### Competing Interests

The authors declare that they have no significant competing financial, professional or personal interests that might have influenced the performance or presentation of the work described in this manuscript.

### Additional Files

Additional File 1: Supplementary results and figures (.docx)

Additional File 2: Primer, sequence and deep sequencing data (.xlsx)

Additional File 3: Cloning and plasmid sequence information (.docx)

Additional File 4: Statistics (.xlsx)

